# Endothelial stiffening induced by CD36-mediated lipid uptake leads to endothelial barrier disruption and contributes to atherosclerotic lesions

**DOI:** 10.1101/2023.11.08.566338

**Authors:** Victor Aguilar, Elizabeth Le Master, Amit Paul, Sang Joon Ahn, Dana Lazarko, Maria Febbraio, James Lee, Irena Levitan

## Abstract

**Background:** To determine the impact of endothelial stiffening induced by CD36-mediated lipid uptake in the disruption of aortic endothelial barrier and development of atherosclerosis in mouse models of obesity and hypercholesterolemia.

**Approach and Results:** Endothelial-specific inducible downregulation of CD36 results in abrogating the stiffening of aortic endothelium induced by a short-term (6-8 weeks) high-fat Western diet in intact freshly isolated mouse aortas of Cdh5.CreER^T2^CD36^fl/fl^ mice, as assessed by atomic force microscopy. No effect was observed on the stiffness of aortic vascular wall assessed in the same groups of mice by echocardiography. Prevention of WD-induced endothelial stiffening by the downregulation of endothelial CD36 was associated with a protective effect against endothelial barrier disruption, assessed by morphological analysis of VE-cadherin junctions and penetration of Evans blue dye into the aortic wall. These protective effects were independent of the changes in the serum lipid profiles. Furthermore, endothelial specific downregulation of CD36 in hypercholesterolemic Cdh5.CreER^T2^CD36^fl/fl^LDLR^-/-^ mice also led to significant decrease in endothelial stiffening after 4-5 months of high fat diet and a significant decrease in the areas of atherosclerotic lesion. In both models, significant endothelial stiffening was observed specifically in male mice, while female mice exhibited less endothelial stiffening and less severe atherosclerosic phenotype, consistent with endothelial stiffening playing an important role in aortic vascular disease in a sex-dependent way. Mechanistically, we show *in vitro* that CD36-mdiated uptake of long chain saturated fatty acids, particularly palmitic acid, induces endothelial stiffening via activation of RhoA/ROCK pathway. Moreover, palmitic acid-induced endothelial stiffening critically depends on the expression of a RhoA inhibitory protein, Rho-GDI-1.

**Conclusions:** We conclude that stiffening of the aortic endothelium by CD36-mediated uptake of fatty acids contributes significantly to WD-induced vascular dysfunction and atherosclerosis. We further propose that fatty acids may activate RhoA by inducing its dissociation from Rho-GDI-1.

## INTRODUCTION

Multiple studies established that a strong predictor for cardiovascular disease is an increase in the rigidity of arterial blood vessels, or arterial stiffness^1,2^. The detrimental effects of arterial rigidity are attributed to the negative effect of sub-endothelial stiffening on the integrity of the endothelial barrier^1,3^, as it is known that growing endothelial cells on stiff substrates results in cell stiffening and barrier disruption^1,3,4^ Our earlier studies discovered that endothelial stiffening can be induced independently of the arterial rigidity by the uptake of oxidized modifications of LDL, oxLDL^5,6^. We also found that feeding mice Western diet (WD) results in the stiffening of the endothelial monolayer in intact aortas *in vivo*, abrogated by the global deletion of CD36 scavenger receptor^5^, known to mediate oxLDL uptake^7^. The goal of this study is to evaluate the impact of CD36/lipid uptake-mediated endothelial stiffening on aortic vascular dysfunction focusing on endothelial barrier integrity, atherosclerosis development and elucidate further mechanistic insights of lipid-induced endothelial stiffening.

Disruption of the integrity of the endothelial barrier is well known to be key in the early stages of vascular inflammation and atherosclerosis. Multiple pro-inflammatory agents were shown to induce barrier disruption via activation of RhoA/ROCK cascade, formation of stress fibers and increase in endothelial contractility^8^. Earlier studies also showed that endothelial barrier is disrupted by exposing endothelial cells to oxidative modifications of LDL (oxLDL) *in vitro*^9^. We hypothesize, therefore, that lipid/CD36-mediated endothelial stiffening should have a significant detrimental effect on endothelial integrity *in vivo*. To test this hypothesis, we determine in this study whether preventing endothelial stiffening by endothelial-specific deletion of CD36 is barrier protective.

Our earlier studies also established that endothelium of the inner curvature of aorta (aortic arch) is stiffer than endothelium of the thoracic (descending) aorta^5^, the arch region well-known to be susceptible to the development of atherosclerosis due to its exposure to pro-atherogenic disturbed flow^10,11^. An increase in endothelial stiffness in aortic arch was exacerbated by feeding mice WD and abrogated by genetic deficiency of CD36. We proposed, therefore, that lipid-mediated endothelial stiffening of aortic arch contributes to the susceptibility of the region to plaque development. Our previous study, however, was performed in wild type mice fed high fat diet, a model that develops obesity but not atherosclerosis. Here, therefore, we extend our studies to a well-established mouse model of hypercholesterolemia, genetic deficiency of LDL receptor (LDLR), known to develop atherosclerosis^12^ and test whether alleviating endothelial stiffening confers an atheroprotective effect.

Mechanistically, our previous studies of endothelial stiffening focused on oxLDL and its oxidized lipids components^5,13,14^, which induces EC stiffening due to contractile responses via the activation of RhoA/ROCK/MLC_2_ pathway^14^, known to lead to the formation of stress fibers, which may destabilize the endothelial barrier by pulling on the junctional proteins^15,16^. However, it is well known that CD36 not only binds and internalizes oxLDL but also plays a key role in the uptake of long chain fatty acids (LCFAs)^17–19^. Indeed, global loss of CD36 results in a decreased uptake of FAs in multiple tissues including adipose^20,21^. Furthermore, endothelial-specific loss of CD36 is sufficient to decrease incorporation of triglycerides^18,19^. Here, we establish the role of CD36-mediated uptake of LCFAs in endothelial stiffening and elucidate the signaling basis of this effect.

## METHODS

Please see the Major Resources Table in the Supplemental Materials

### Animal and Cellular Models

All studies involving animals were approved by the Institutional Animal Care and Use Committee at the University of Illinois at Chicago. To generate an endothelial-specific deficiency of CD36, floxed CD36 mice (CD36^fl/fl^, gift of Prof Maria Febbraio) in the C57BL/6J background were utilized. These mice were generated to target exons 2 and 3, sites containing the translation start site and the first transmembrane domain^22^. These mice were crossed with mice with a VE-cadherin promoter Cre.ER^T2^ (Cdh5.CreER^T2^;CD36^fl/fl^) which have inactive Cre recombinase until delivery of tamoxifen. These mice were also crossed with mTmG reporter mice (Cdh5.CreER^T2^;mTmG;CD36^fl/fl^) to track endothelial specificity. A second strain was developed by crossing Cdh5.CreER^T2^CD36^fl/fl^ with LDLR^-/-^ mice to produce Cdh5.CreER^T2^;CD36^fl/fl^;LDLR^-/-^ mice, suitable for the development of atherosclerosis. Animals are fed commercial (TD.88317, from Inotiv) Western-type diet (21% weight from milk, 34.5% weight from sucrose) or standard low-fat diet. Mice were either euthanized employing CO_2_ or isoflurane, followed by cervical dislocation or pneumothorax (decapitation used for collection of blood for serum analysis). Aortas were isolated for clean-up and experimental analyses.

All *in vitro* work was performed with low passage HAECs (passage 5-8)^5^.

### Analysis of Fasted Serum Lipids

Blood was collected following a 6 hour fast. Following euthanasia, blood was collected for serum extraction *via* centrifugation. Samples were assessed for lipid panels consisting of low-density lipoprotein, high-density lipoprotein, total cholesterol, triglycerides, and glucose.

### Immunostaining, Quantification of Vascular Leakage

Immunostaining was performed on freshly-isolated aortas using standard protocol. A whole-mouse cardiac perfusion/fixation was performed through the left ventricle by perfusing with heparin saline solution via a syringe pump, followed by a 2% paraformaldehyde. Fixed tissues were permeabilized for an hour at room temperature (RT) with 0.2% Triton X-100 and blocked for an hour at RT with 5.5% donkey serum. Primary antibodies for anti-mouse CD36 (Biotechne, af2519) and anti-mouse VE-cadherin (BD Biosciences, 555289) were added and incubated overnight at 4°C. Secondary antibodies for anti-goat Alexa Fluor 488 (Thermo-Fisher, A21206) and anti-rat Alexa Fluor Cy3 (Jackson, 712-165-153) were added along with DAPI (Thermo-Fisher, D1306) and incubated for 3 hrs at RT. Images were taken using a Zeiss LMS 710/META confocal microscope, using 63X/oil magnification. VE-cadherin width was quantified using a machine vision algorithm program. Microarchitecture of the junctions employs a customized segmentation algorithm that generates isosurfaces based on fluorescence intensity values. Thickness is defined and measured perpendicular to these segments in voxels. Voxels are converted to microns to assign diameter values to these segments. EBD was injected in retro-orbital sinus to assess barrier disruption. After circulation, aorta images are acquired with fluorescence confocal microscopy.

### Pulse Waveform Velocities

were measured using echocardiographs using Doppler imaging, and mean PWV was generated from calculating values over 10 cardiac cycles obtained during echocardiography done in each mouse^23^.

### siRNA Transfection, Real Time-Quantitative Polymerase Chain Reaction

HAECs were transfected with siRNA designed by Qiagen to target Rho-GDI-1 and the downregulation of mRNA expression confirmed by PCR using a standard protocols. PCR was also performed on freshly-isolated lung endothelial cells, as described earlier.

### Oil Red O Staining

In select mice, aortas were prepared for Oil Red O staining towards the quantification of atherosclerotic lesion area. Visualization was done by staining on Oil Red O solution for 30 minutes, followed by washing in an isopropanol:DI water solution. Pictures were captured with a camera mounted on a brightfield microscope, always taken with a metric ruler. Quantification of the lesion area was done blind using ImageJ software.

### Atomic Force Microscopy

Elastic modulus values of cells and aortas were measured with an Asylum MFP-3D-Bio atomic force microscope. Gold-coated silicon nitride cantilevers (typical spring constant of 0.1 N/m) were employed exclusively for these experiments. We employed a descent velocity of 2 µm/s and a trigger force of 5-10 nN. Based on these values, the general indentation into the cell was approx. 15% of the total cell height. Force-distance curves were analyzed employing the Hertz model as described in prior publications^5^. The experimental curves were fitted into bi-domain polynomial models using a standard least-squares minimization algorithm. For cultured cells, measurements were obtained from the region between the periphery and the nucleus. For aortic tissues, samples were first excised from aortas. Excisions were done as denoted in our experiments from either atherogenic regions (e.g. the inner aortic arch) or atheroprotective regions (e.g. the descending aorta). After their excision, samples were allowed to stick onto double-sided tape to ensure attachment. 15 to 20 measurements per sample were gathered.

### Statistical Analyses

Results are expressed as the mean ± SEM or individual values arrayed into histograms/dot plots. Sample sizes were determined through power analysis and equality of variance between groups assessed via F-test. Significant differences were determined by a Student’s t-test or an ANOVA multiple comparison test. A p-value of less than 0.05 was considered significant.

## RESULTS

### Endothelial-specific downregulation of CD36 protects aortic endothelium against WD-Induced stiffening

#### Generation and validation of inducible endothelial-specific downregulation of CD36

CD36^fl/fl^ mice were crossed with a tamoxifen-inducible endothelial specific *Cdh5*.CreER^T2^ mouse strain, also bred to contain a ROSA^mTmG^ reporter to generate Cdh5.CreER^T2^;mTmG;CD36^fl/fl^ that allows efficient identification of successful Cre activation/CD36 deletion^24^. Endothelial-specific deletion of CD36 was induced at 8 weeks of age and verified by RT-qPCR in endothelial cells freshly-isolated from the lungs and by CD36 immunostaining performed on intact aortas *ex vivo*. RT-qPCR showed that expression of CD36 in endothelial cells, identified by an endothelial marker CD31, was decreased by ∼80% whereas no decrease was observed in vascular smooth muscle cells, identified by a VSMC marker, *Acta2* (*<u>Fig.1Aa</u>*). As expected, injection of tamoxifen efficiently turned the mTmG reporter from red to green in the endothelium monolayer (*<u>Fig.1Ab</u>*, upper panels) and significantly decreased the expression of CD36 in aortic endothelium (*<u>Fig.1Ab</u>* lower panels, *<u>Fig.1Ac</u>*), as visualized by confocal microscopy. Fluorescence imaging in recent studies has demonstrated that CD36 localizes to both the nuclear and the cytoplasmic areas of the cell^18,25^. These confocal microscopy results show a similar distribution of CD36.

#### Downregulation of endothelial CD36 prevents WD-induced stiffening of aortic endothelium in male mice

To determine the impact of CD36 deletion on WD-induced endothelial stiffening, Cdh5.CreER^T2^CD36^fl/fl^ mice injected either with tamoxifen (labeled EC-CD36^-/-^) or with vehicle (labeled EC-CD36^WT/WT^) were put either on WD (21% weight from fat) or maintained on low fat diet (LFD) for 6-8 weeks. We also employed Cdh5.CreER^T2^ mice injected with tamoxifen or vehicle and fed WD as additional controls to exclude the off-target effects of tamoxifen. As expected, this short-term WD resulted in an increase in weight and elevated levels of fasting total cholesterol, LDL and HDL, but the downregulation of endothelial CD36 had no effect on either of these parameters (*<u>Fig.S1A,a-d</u>*). No changes were observed in triglycerides and glucose (*<u>Fig.S1A,e-f</u>*). The stiffness of the endothelium (elastic modulus, EM) of intact freshly isolated aortas was measured using AFM. In EC-CD36^WT/WT^ male mice, consistent with our previous studies, WD elicited a substantial increase in the elastic modulus of aortic endothelium in freshly excised aortas, as compared to the endothelium of aortas excised from mice maintained on LFD, indicative of EC stiffening, as is evident from a shift in the histogram distribution of EM values from left to right (*Fig.1B<u>, top row</u>*) and the distribution of the average EM values in individual mice (*Fig.1C<u>, left)</u>*. Endothelial-specific downregulation of CD36 (EC-CD36^-/-^) prevented WD-induced EC stiffening pattern (*Fig.1B<u>, bottom row</u>*) and the average EM values in individual mice (*Fig.1C<u>, right</u>*). AFM experiments performed on age-matched Cdh5.CreER^T2^ controls showed that tamoxifen injections do not affect WD-induced EC stiffening (*<u>Fig.S2A,a-c</u>*) or serum lipid panel (*<u>Fig.S2B,a-e</u>*).

**Figure 1.**
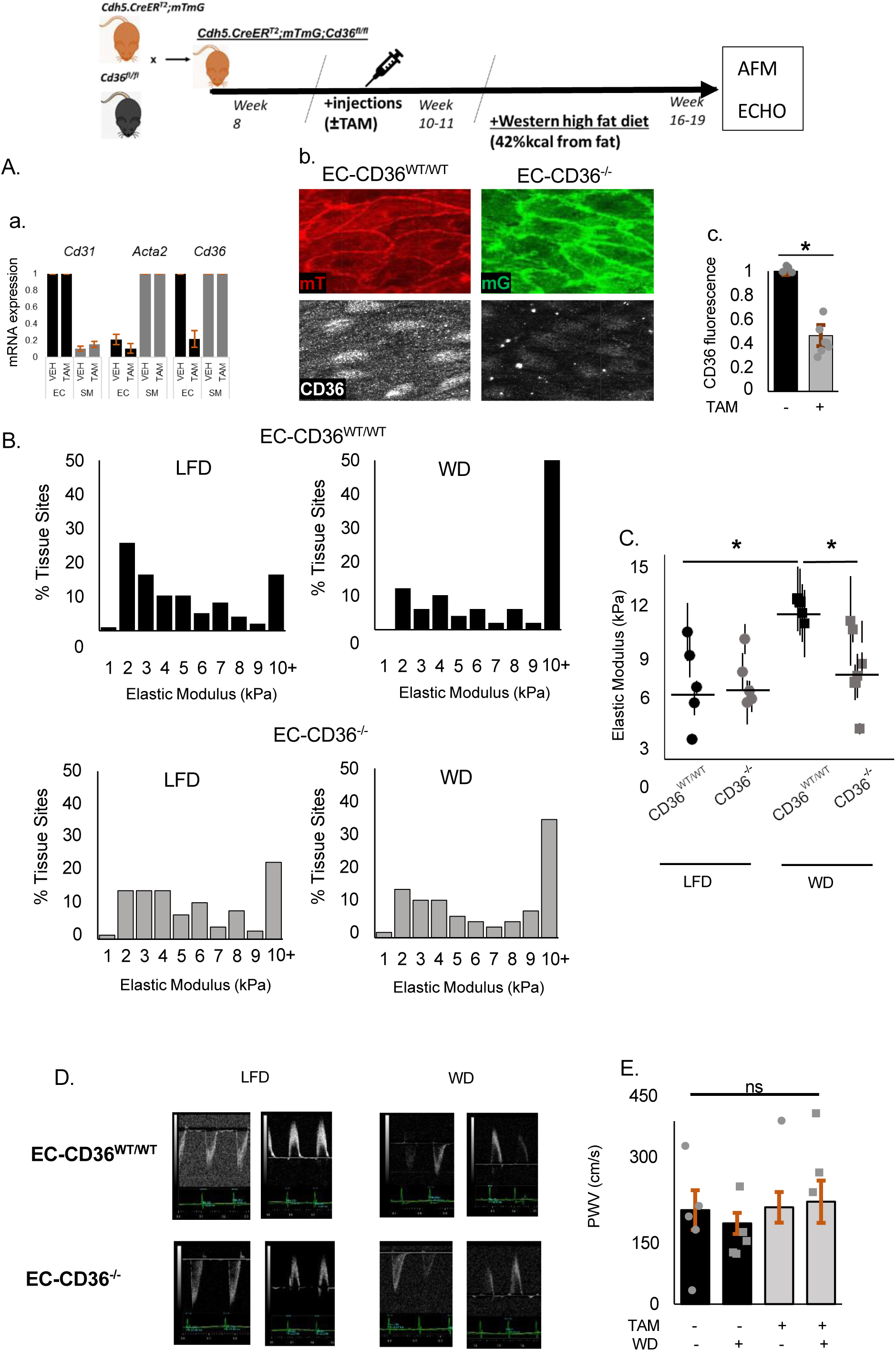
Endothelial-specific deficiency of CD36 prevents short-term WD-induced stiffening of aortic endothelium in male mice. Pictoral diagram describing the timeline for the short-term (6-8 week) WD regimen. **A)** *<u>Validation of the endothelial-specific deficiency of CD36 in the mouse model</u>*: **a)** RT-PCR done on isolated lung ECs to assess mRNA expression of CD36 in Cdh5.CreER^T2^;mTmG;CD36^fl/fl^ injected with vehicle (VEH) or tamoxifen (TAM); CD31 and Acta2 are used as endothelial and smooth muscle markers respectively (n=3 mice pairs). **b)** CD36 immunostaining in aortas of the same mice mice: the upper panels show mTmG Cre-sensitive reporter in mice injected with the vehicle (left panel, red) or injected with tamoxifen (right panel, green). The lower panels show CD36-specific imunoflorescence in the aortas of the same mice. **c)** mean CD36 fluorescence (n=7 mice pairs). **B)** Histograms of elastic moduli of aortic endothelium measured in mice of 4 experimental groups: EC-CD36^WT/WT^mice (injected with vehicle) or EC-CD36^-/-^ mice (injected with tamoxifen), both on LFD or WD (5-6 mice per group at 10-15 distinct sites for each aorta (pooled data)). **C)** Mean elastic moduli for individual mice in the same experimental groups as B (p-values were calculated by one-way ANOVA). **D)** Representative Doppler waveforms (left = ascending, right = descending) of echocardiograms of aortas in the same experimental groups and **E)** Mean pulse waveform velocities (n=5 mice per group).

#### WD-induced stiffening was specific to the endothelium

In contrast to endothelial stiffening, the short term WD used in this study did not affect arterial stiffness evaluated through pulse wave velocity (PWV) measurements, a method commonly used to evaluate the stiffness of the arterial vascular wall^23^. Endothelial downregulation of CD36 had no effect on arterial stiffness neither on WD nor on LFD (*Fig.1D<u>,E</u>*).

#### WD-induced endothelial stiffening was not observed in female mice

To compare endothelial stiffness in male and female mice, we first examined aortas harvested on the same day from age-match male and female mice maintained on low fat diet. No difference was observed in the elastic moduli of the endothelium between males and females (*<u>Fig.S3A,a-c</u>*). However, the impact of WD on endothelial stiffness in females was significantly different from males: in contrast to males, a separate cohort of age-matched female mice maintained on WD for the same period of time, as males, had only a small and not statistically significant increase the endothelial elastic modulus and consequently no significant decrease in stiffness upon deletion of endothelial CD36 (*<u>Fig.S3B,C</u>*). Female mice also had less WD-induced weight gain and lower levels of serum cholesterol than their male counterparts (*<u>Fig.S1B,a-f</u>*).

### Endothelial-specific downregulation of CD36 protects against WD-Induced barrier disruption in aortic endothelium

To evaluate the functional significance of CD36-mediated stiffening of aortic endothelium, we assessed the impact of CD36 on the integrity of the endothelial barrier in intact aortas by the morphological analysis of adherens junctions, a key determinant of the barrier integrity^26^, followed by a functional analysis of endothelial leakage.

#### Morphological analysis of adherens junctions

was performed by immunostaining for VE-cadherin, confocal imaging and machine-vision analysis of the intact endothelial layer of aortas freshly isolated from the same experimental groups of mice, as described above: Cdh5.CreER^T2^CD36^fl/fl^ mice injected with tamoxifen or vehicle maintained on WD or LFD diet for 6-8 weeks (*Fig.2A*). CD36 immunostaining was performed on the same samples. As expected, VE-cadherin junctional morphology altered from smooth and thin in aortas of animals maintained on LFD to wide and jagged in response to WD, which is indicative of disruption of the barrier^3,27^. Downregulation of CD36 expression had no effect on the junctional width in animals maintained on LFD but had a strong protective effect against the widening of the junctions in animals on WD. The widening of junctions on WD and the protective effect of the loss of CD36 are apparent in the representative images of VE-cadherin (2A, the upper row) and verified with machine vision showing the color-coded measurements of VE-cadherin junctional A representative set of videos displays 3D reconstructions of the confocal microscopy data (and accompanying machine vision of the junctional width) from the aorta scans for EC-CD36^WT/WT^ mice on WD (*<u>Videos.S1-3</u>*) versus EC-CD36^-/-^ mice on WD (*<u>Videos.S4-6</u>*).

**Figure 2.**
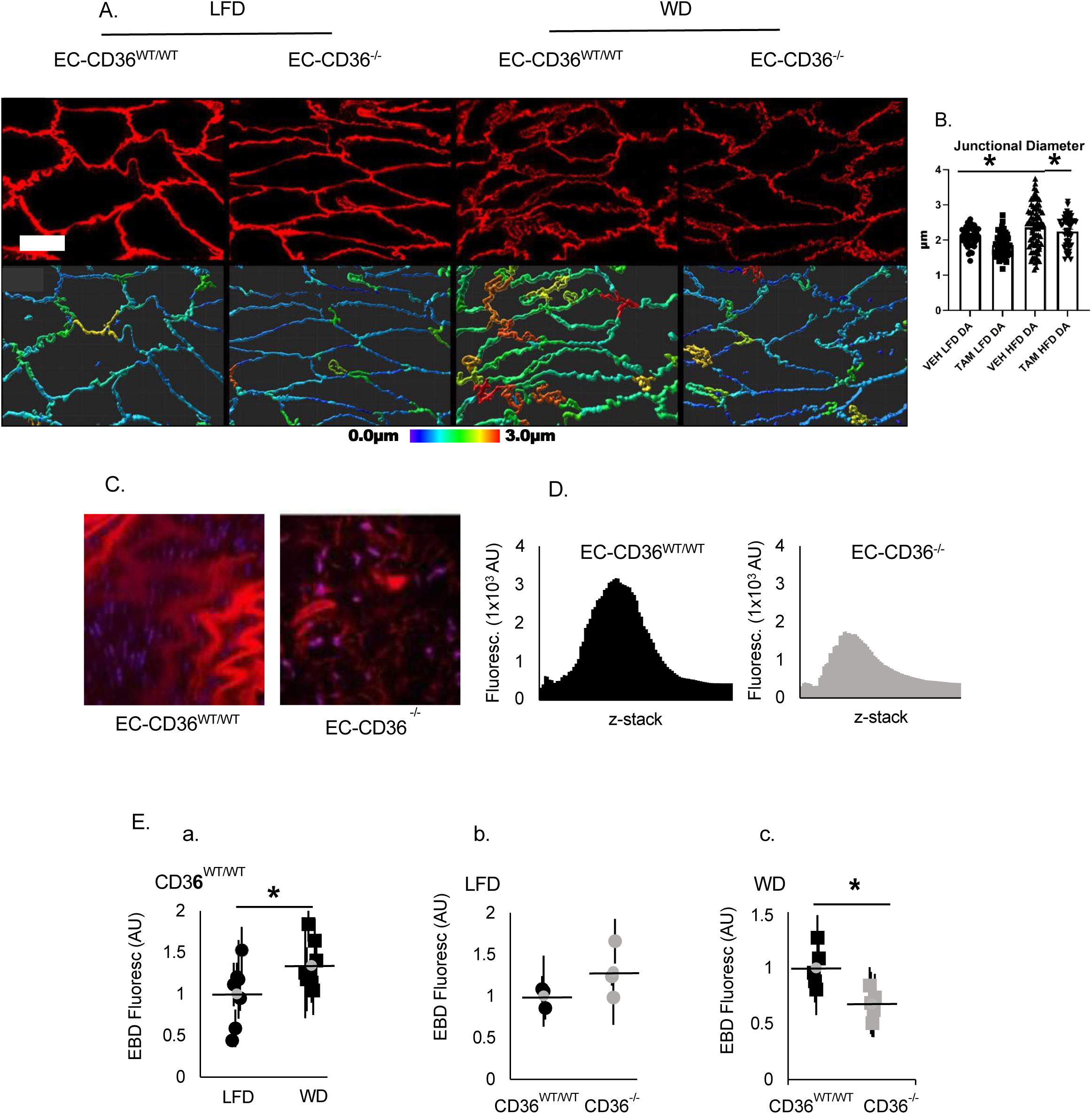
Endothelial-specific deficiency of CD36 protects against short-term WD-induced disruption of the endothelial barrier. **A)** VE-cadherin immunostaining of *en face* endothelium of thoracic aortas in the four experimental mice groups defined in figure 1: representative VE-cadherin fluorescence (top row) and same images color-coded for the junctional width (bottom row). **B)** Quantification of VE-cadherin width in the same representative mice cohort (n=3-4 cohorts analyzed). **C)** Representative confocal images of EBD in excised aortas 1 hour following retro-orbital injection of dye to mice and **D)** corresponding z-stack scan of the EBD fluorescence intensity showing the penetration of EBD into the aorta. **E)** Quantification of aortic mean EBD fluorescence intensity in three separate cohorts: **a)** EC-CD36^WT/WT^ on LFD vs WD (n=7-8 mice ger group), **b)** EC-CD36^WT/WT^ vs EC-CD36^-/-^ controls (LFD-fed n=3 mice per group), **c)** EC-CD36^WT/WT^ vs EC-CD36^-/-^ (WD-fed n=5 mice per group).

#### Endothelial CD36 deficiency reduces endothelial leakage resulting from WD

Endothelial leakage was quantified using Evans blue dye (EBD) that leaks into the vascular wall in areas of barrier disruption. Since EBD fluoresces when bound to albumin, the penetration of the dye into the tissues was tracked by confocal microscopy (*Fig.2C<u>, left</u>*). Consistent with the protective effect of the downregulation of CD36 on the width of the VE-cadherin junctions, it also resulted in a significant decrease in the EBD leakage in mice fed WD (see representative single section images in 2C and histograms of the corresponding z-stacks, 2D). Total EBD fluorescence in aortas of individual mice show that WD resulted in an increase in barrier disruption, as expected (*Fig.2E<u>,a</u>*) and that while CD36 deficiency has no effect on the barrier in mice on LFD (*Fig.2E<u>,b</u>*), it has a strong protective effect against barrier disruption induced by WD (*Fig.2E<u>,c</u>*). No effect of tamoxifen was observed in Cre controls on WD (*<u>Fig.S4A</u>*). Tracking the depth of EBD penetration into the aorta showed no differences across all experimental groups studied (*<u>Fig.S4B</u>*).

### Endothelial stiffening correlates with atherosclerotic lesion formation in male and female LDLR^-/-^ mice

#### Increase in endothelial stiffness in aortas of LDLR^-/-^ male mice

In order to determine if there was a correlation between endothelial stiffening and atherosclerotic plaque formation, we employed a well-known hypercholesterolemic LDLR^-/-^, since mice with a WT B6 background do not develop atherosclerosis. Focusing first on male mice, we show that aortic endothelial monolayer of LDLR^-/-^ mice is significantly stiffer (higher elastic moduli) than that of the WT controls, both at athero-resistant (DA) and athero-susceptible (AA) regions, as apparent from a rightward shifts in the elastic modulus histograms (*Fig.3A<u>: compare upper and lower rows</u>*) and the mean values for individual mice (*Fig.3B*). The experiments are performed in 6-7 month old mice. Furthermore, as we previously showed in WT mice^5^, we now show that the endothelium of the pro-atherogenic arch is significantly stiffer than endothelium in the athero-protective descending aorta in the LDLR^-/-^ mouse (*Fig.3A<u>: compare left and right columns,</u> Fig.3B*). The values of the elastic moduli of the AA endothelium of LDLR^-/-^ mice, which are around 25 kPa are also significantly higher than a typical range between 5 and 15 kPa in aortas of WT mice^5^.

**Figure 3:**
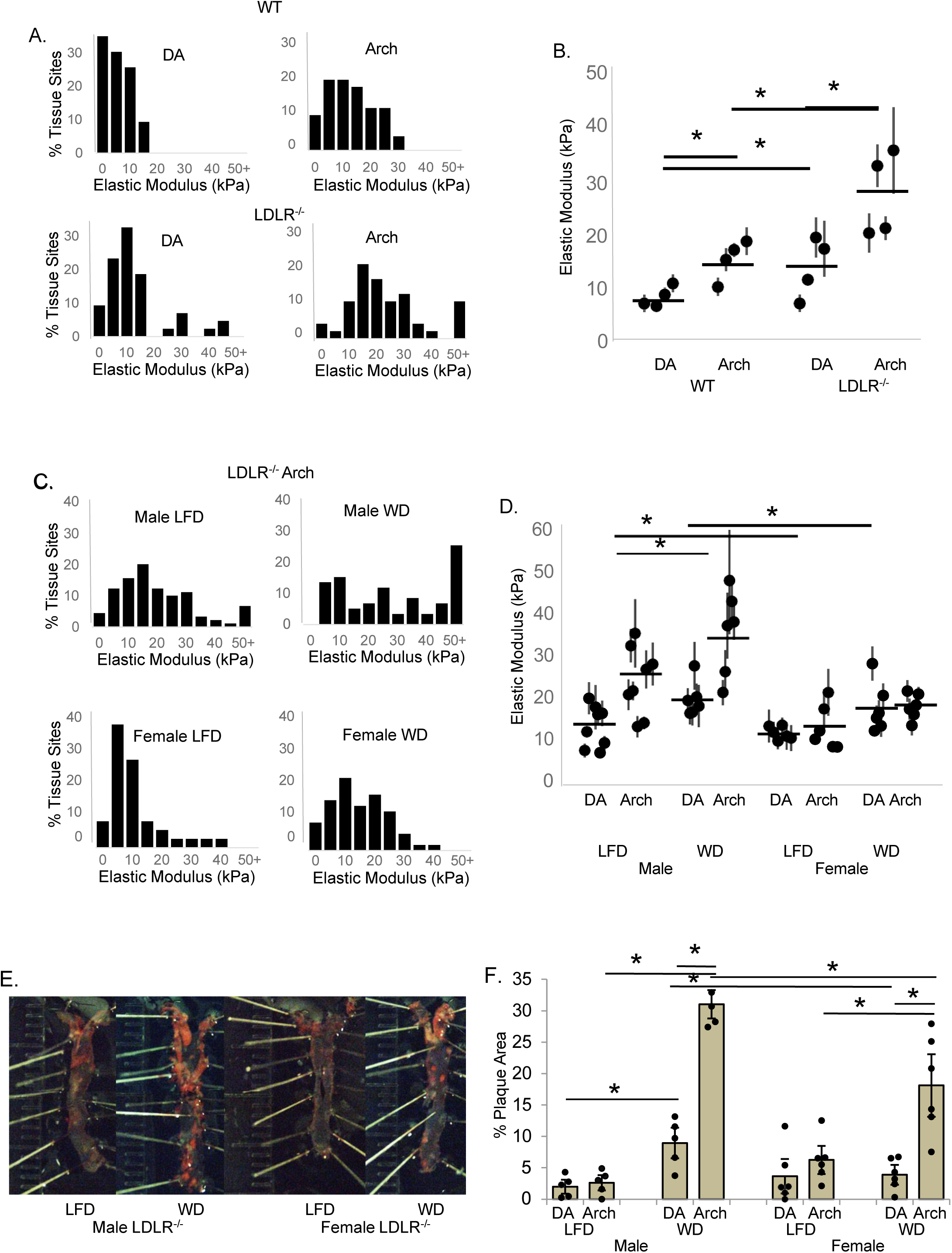
Hyperlipidemia resulting from the loss of the LDL-receptor increases endothelial stiffness especially in the aortic arch of male mice with a corresponding increase in aortic plaque area. (**A**) Histograms and **(B)** averaged elastic modulus values for the DA (descending aorta) and AA (aortic arch) in WT versus LDLR^-/-^ male mice (n=4, 10-20 sites/sample). **(C)** Histograms and **(D)** averaged elastic modulus values for male and female LDLR^-/-^ mice on low fat diet or 4-5 month western diet (n=5-8, 10-20 tissue sites/sample). **(E)** Representative images of Oil O Red stained aortic lipid deposition and **(F)** averaged percent plaque coverage quantified in the aortic arch and descending aorta (abdominal and thoractic aorta combined). N=5-6 mice/condition.

#### Differential effects of WD on endothelial stiffness in male and female LDLR^-/-^ mice

It is also known, however, that while LDLR^-/-^ mice are hyperlipidemic genetically, atherosclerotic lesions develop spontaneously only at later ages, a process strongly exacerbated by WD^28^. In this series of experiments, cohorts of male or female LDLR^-/-^ littermates were fed WD or LFD for 4-5 months, a period chosen to allow time for the development of atherosclerotic plaques. AFM measurements of aortic endothelial stiffness were performed simultaneously on age-matched male and female LDLR^-/-^ mice in the DA and AA, on LFD and WD. In *male* LDLR^-/-^ mice WD resulted in significant endothelial stiffening in both DA and AA areas of the aorta, as compared to LFD, with an enhanced effect observed in the AA (*Fig. 3C top row, D*). In *female* LDLR^-/-^ mice, however, no significant endothelial stiffening developed in response to WD and no differential stiffness was observed between DA and AA regions (*Fig. 3C bottom row, D*). These results highlight the fact that endothelial stiffness from intact aortas is sex-, regionally-, and diet-dependent in LDLR^-/-^ mice.

#### Correlation between endothelial stiffening and the severity of plaque formation in male vs

*female LDLR^-/-^ mice*: Plaque areas were analyzed using Oil Red O *en face* staining for lipids in the entire aorta, including the aortic arch and the descending aorta (thoracic and abdominal aortic segments) isolated from male and female LDLR^-/-^ mice either fed a WD for 4-5 months or maintained on the regular LFD. As expected^28^, these mice when maintained on LFD have only few lesions (1-3% of the area of the aorta in male and 2-10% in female mice), which is significantly increased in mice fed a WD (*Fig.3E<u>,F</u>*). However, a significantly greater plaque deposition was observed in male WD-fed LDLR^-/-^ mice, as compared to age-matched females: in male mice, a plaque area increases to 9 and 31% in DA and AA regions respectively, whereas in female mice, there was no increase in the plaque area in the DA region and an increase only to 18% in the AA. No significant difference between males and females was found in the lipid panel parameters or weight gain (*<u>Fig.S5)</u>*. An increased plaque depositions in male mice vs. females correspond to an increase in endothelial stiffness.

### Endothelial-specific downregulation of CD36 abrogates WD-induced endothelial stiffening in LDLR^-/-^ mice and confers a protective effect against plaque development

#### Role of endothelial CD36 in endothelial stiffening in hypercholesterolemic LDLR^-/-^ mice

was tested by cross-breeding Cdh5.CreER^T2^CD36^fl/fl^ mice with LDLR^-/-^ mice to generate inducible EC-specific CD36 deficiency model on hypercholesterolemic LDLR^-/-^ background. AFM experiments were performed on Cdh5.CreER^T2^CD36^fl/fl^LDLR^-/-^ sets of male littermates, injected either with tamoxifen or vehicle control at 8 weeks of age, as described above for Cdh5.CreER^T2^CD36^fl/fl^ mice, and then maintained on WD or LFD for 4-5 months. Cdh5.CreER^T2^CD36^fl/fl^LDLR^-/-^ mice injected with vehicle control (EC-CD36^WT/WT^;LDLR^-/-^) developed the same pattern of endothelial stiffening, as we observed in LDLR^-/-^ mice: higher elastic moduli in the AA region, as compared to the DA, and increased stiffness in response to WD, particularly in the AA region (*Fig.4A*, top panels, B, left, histograms shown for AA regions only). Downregulation of endothelial CD36 by tamoxifen (EC-CD36^-/-^LDLR^-/-^ mice) has a profound effect on this pattern: no difference was observed in the elastic moduli between the AA and DA regions and no endothelial stiffening developed in response to WD (*Fig.4A*, bottom panels, B, right). Overall, the values of endothelial elastic modulus in EC-CD36^-/-^LDLR^-/-^ mice are lower than in EC-CD36^WT/WT^LDLR^-/-^ mice across all the experimental groups of animals described here: LFD vs WD and DA vs AA. Importantly, no tamoxifen effect on endothelial stiffness was observed in control Cdh5.CreER^T2^ mice (*<u>Fig.S6</u>*). These data indicate that endothelial CD36 plays a crucial role in the control of aortic endothelial stiffness in hypercholesterolemia.

**Figure 4:**
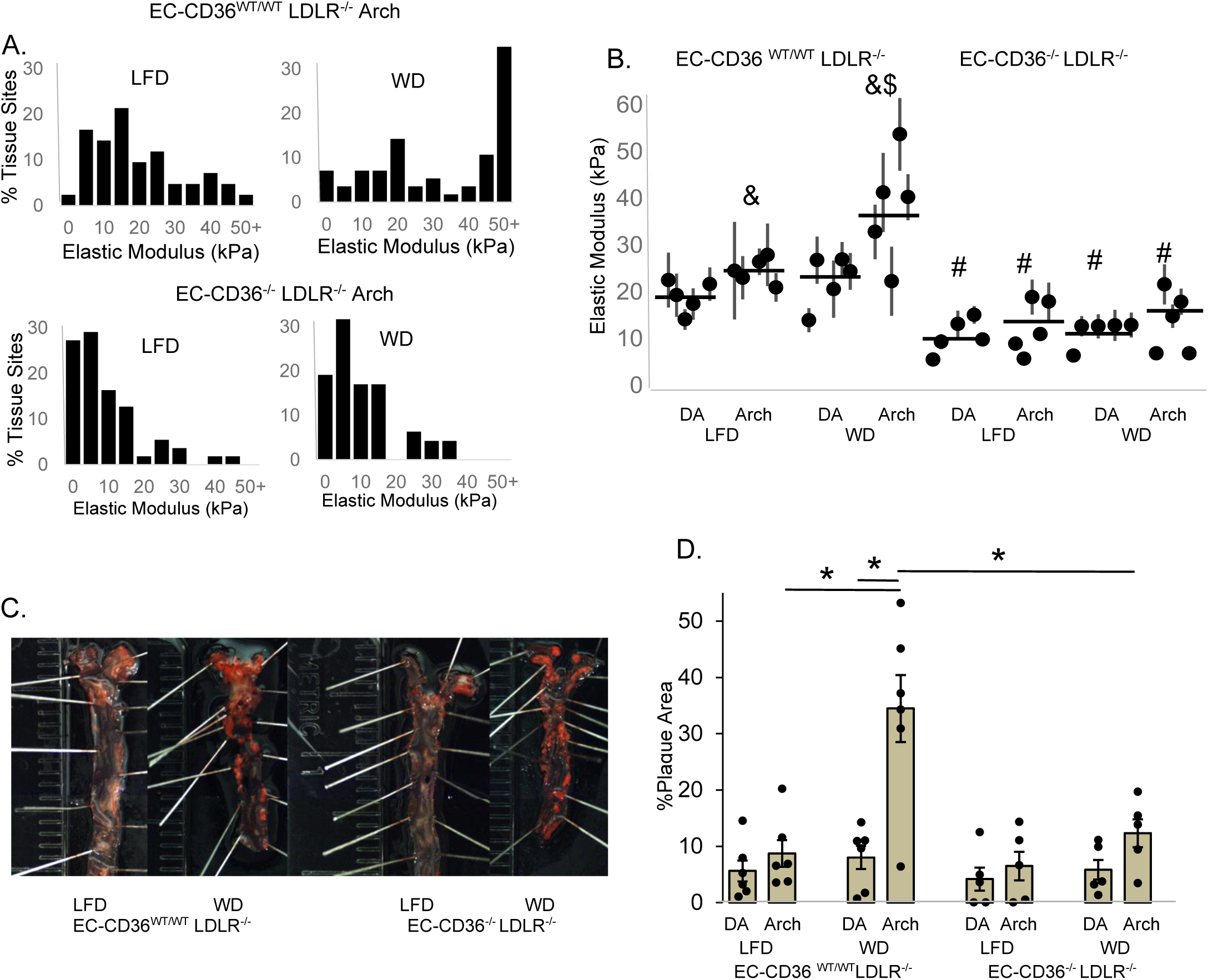
Endothelial stiffening in the aortic arch and percent plaque area is dependent on endothelial CD36 in a 4-5 month high fat WD feeding. (**A**) Histograms and **(B)** averaged elastic modulus values of EC-CD36^WT/WT^LDLR^-/-^ (vehicle) controls compared to EC-CD36^-/-^LDLR^-/-^ (tamoxifen-injected) male mice (n=5, 10-20 tissue sites/sample) in the DA and AA in LFD and WD-fed mice. **(C)** Representative images of Oil Red O stained aortic lipid deposition and **(D)** averaged percent plaque area in the DA and AA in LFD and WD-fed male control EC-CD36^WT/WT^LDLR^-/-^ versus EC-CD36^-/-^LDLR^-/-^mice (n=5-6). *p<0.05. &p<0.05 as compared to its DA counterpart #p<0.05 as compared to its vehicle control counterpart $p<0.05 as compared to its LFD counterpart

#### Role of endothelial CD36 in plaque formation in LDLR^-/-^ mice

We next show that prevention of WD-induced endothelial stiffening by the downregulation of endothelial CD36 strongly correlates with a decrease in the atherosclerotic burden of the aorta, as assessed by *en face* Oil Red O staining (*Fig.4C<u>,D</u>*). As shown in the representative images in Fig. 4C there is significantly less plaque area in aortas of EC-CD36^-/-^LDLR^-/-^ mice as compared to EC-CD36^WT/WT^ LDLR^-/-^ mice. Indeed, similarly to LDLR^-/-^ mice, EC-CD36^WT/WT^LDLR^-/-^ mice fed WD developed significant plaque depositions in the AA region with 35% of the plaque area, a significant increase from 8% observed in aortas of mice maintained on LFD. In contrast, EC-CD36^-/-^LDLR^-/-^ mice fed WD had no significant increase in plaque deposition neither in the DA, nor in the AA areas indicating a strong protective effect. In terms of plasma lipids, downregulation of endothelial CD36 had no significant effect on the levels plasma lipoproteins, LDL and HDL, elevated as expected by WD, but attenuated the elevation of the total cholesterol and triglycerides level (*<u>Fig.S7</u>*).

### Saturated LCFAs induce EC stiffening in vitro and ex vivo

#### Differential effects of LCFAs on endothelial stiffening of HAECs

Since CD36 is a receptor for LCFAs, which also have pro-inflammatory effects^29–31^, we next tested the impact of 3 common LCFAs, two saturated LCFA, palmitic and stearic, and one unsaturated, oleic, on HAEC stiffness. The histograms of the elastic moduli show the combined data for all the experiments (*Fig.5A*) and the plots show the means for the individual samples (12-20 cells measured in every sample) (*Fig.5B*). This experiment showed that exposure to saturated but not to unsaturated FAs results in significant EC stiffening, with the strongest effect observed for palmitic acid (*Fig.5A<u>,B</u>*). Furthermore, we also show that palmitic acid-induced stiffening in CD36 dependent: To block CD36-mediated transport, HAECs were exposed to SSO, which covalently crosslinks residue Lys164 in the FA/oxLDL ectodomain, irreversibly blocking CD36-mediated uptake of both lipids^32^. As we have shown previously^6^, SSO prevented oxLDL-induced stiffening of HAECs (*Fig.5D*). Here we show that SSO also prevents palmitic acid-induced HAEC stiffening indicating the role of CD36 (*Fig.5C<u>,D</u>*).

**Figure 5:**
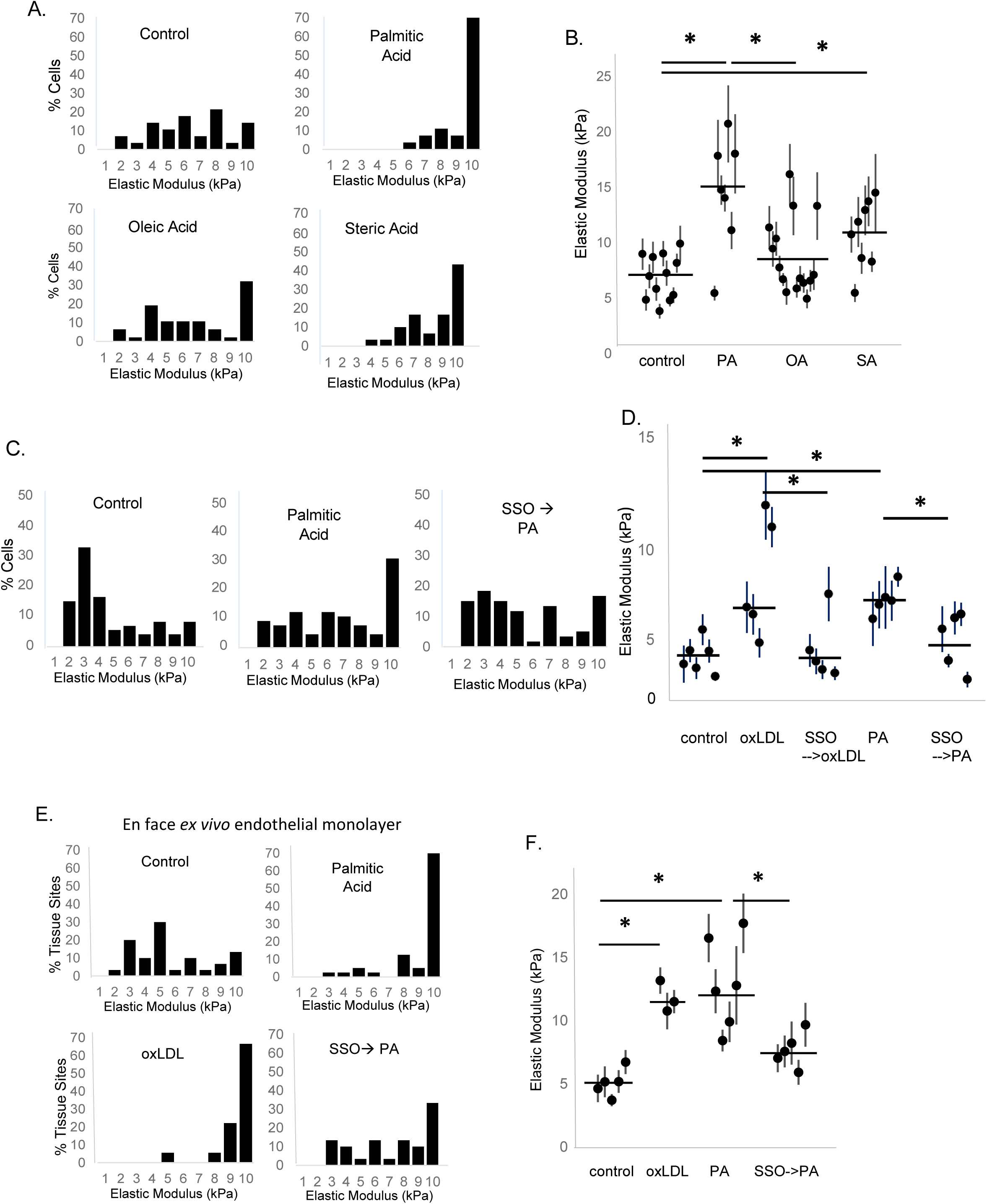
Palmitic acid but not oleic acid induces endothelial stiffening in a CD36 dependent manner *in vitro* and *ex vivo* in intact endothelium in WT controls. (**A**) Histograms and **(B)** average elastic modulus stiffness values in HAECs treated with 5 µM concentration of either palmitic acid, oleic acid, or stearic acid (N=5-12 independent experiments with 12-20 cells sampled per condition per experiment). **(C)** Histograms and **(D)** averaged elastic modulus values in HAECs treated with (10 µg/mL) oxLDL or PA with pre-treatments of the CD36 irreversible inhibitor, SSO (25 µg/mL) (N=5-6 independent experiments with 12-20 cells/condition/experiment). **(E)** Histograms and **(F)** averaged elastic modulus values from the endothelial monolayer of intact descending aortas from WT males with either oxLDL or PA with the pre-treatment SSO (n=4 mice with 8-15 tissue sites per condition). *p<0.05

#### EC stiffening by palmitic acid is observed in aortic endothelium and is dependent on CD36

We have shown previously that exposing intact aortas to oxLDL ex vivo results in significant stiffening of the endothelium monolayer^5^. Here we show that the same effect is induced by exposing aortas freshly-isolated from WT mice to palmitic acid, with oxLDL used as a positive control (*Fig.5D<u>,E</u>*). Furthermore, similarly to the effect in vitro, exposing aortas to SSO abrogates palmitic acid-induced stiffening effect.

### Palmitic acid-induced endothelial stiffening is mediated by Rho signaling via Rho-GDI-1

#### PA-induced EC stiffening is mediated by ROCK but not by Src

Our earlier work showed that oxidized lipids induce EC stiffening through Rho-associated protein kinase (ROCK), which phosphorylates myosin light chain towards the generation of contractile stress fibers^13,14^. We show here that PA-induced endothelial stiffening is also abrogated by inhibiting ROCK with its pharmacological inhibitor Y27632 (*Fig.6A<u>,B</u>*). Baseline stiffness for ECs did not change in response to ROCK inhibition (*Fig.6B*). The role of Src kinases was tested because their known roles in CD36-induced signaling^33^. Specifically, we used PP2, defined as a pan-inhibitor of Src^34^. We show, however, that PP2 has no effect on either oxLDL-induced or PA-induced endothelial stiffening.

**Figure 6:**
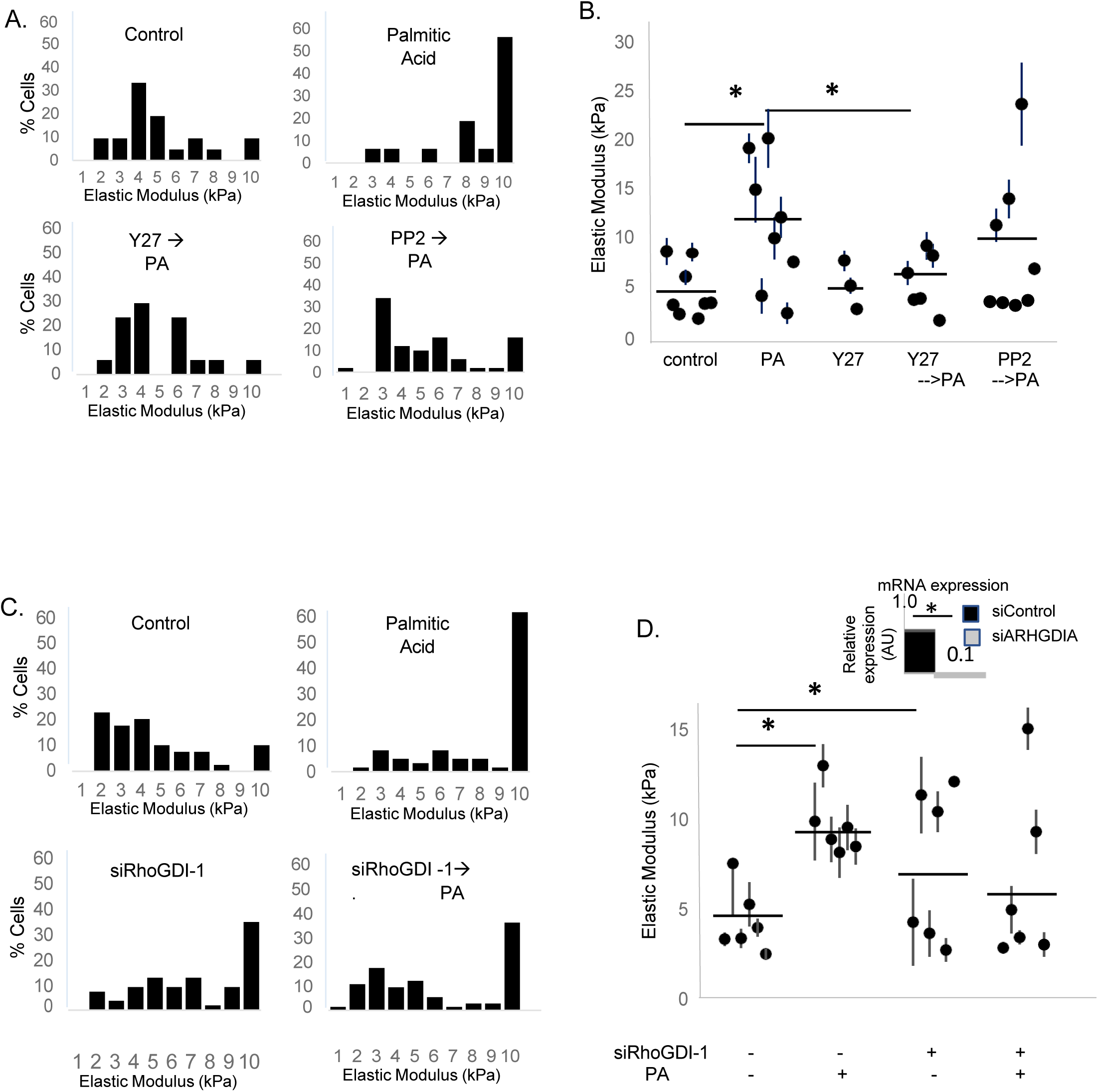
Palmitic acid-induced endothelial stiffening requires Rho protein kinase but not Src kinase and loss of RhoGDI-1 induces endothelial stiffening. (**A**) Histograms and **(B)** average elastic modulus stiffness values in HAECs treated with 5 µM palmitic acid with pre-treatments of Rho kinase inhibitor Y27632 (1 µg/mL) or Src kinase inhibitor PP2 (15 µM) (N=3-8 independent experiments with 12-20 cells sampled/experiment). **(C)** Histograms and **(D)** averaged elastic modulus values in HAECs treated with PA with or without downregulation of RhoGDI-1 (N=6 independent experiments, 8-20 cells per condition/experiment). Inset contains downregulation of RhoGDI-1. *p<0.05

#### PA-induced EC stiffening requires Rho-GDI-1

To elucidate the mechanism of PA activation of RhoA/ROCK cascade, we addressed the hypothesis that it might be mediated by the disruption of an inhibitory interaction between RhoA and Rho-GDI-1, known to associate via a hydrophobic pocket^35^. To test this hypothesis, we downregulated Rho-GDI-1 in HAECs using siRNA resulting in a decrease by >90%, as verified by RT-qPCR (*Fig.6D<u>, inset</u>*). The loss of Rho-GDI-1 *per se* resulted in an increase in the endothelial elastic modulus, as expected because of the removal of its inhibitory effect on RhoA. Most importantly, downregulation of Rho-GDI-1 abrogated the stiffening effect of palmitic acid indicating that it is essential for the effect (*Fig.6C<u>,D</u>*).

## DISCUSSION

Endothelial stiffness is a fundamental biomechanical property of the inner lining of the blood vessels, an interface between the lumen of the vessel and the vascular wall. Our earlier studies discovered that high fat WD increases the stiffness of the endothelium layer in mouse aortas, an effect abrogated by the global deletion of a scavenger receptor CD36, leading us to the hypothesis that lipid-mediated endothelial stiffening may play an important role in WD-induced endothelial dysfunction^5^. Here, we address this hypothesis by selectively abrogating the stiffening of the endothelium by inducible endothelial-specific downregulation of CD36 in WT and in hypercholesterolemic LDLR^-/-^ mice. We demonstrate that the downregulation of endothelial CD36 prevents WD-induced endothelial stiffening in both WT and LDLR^-/-^ mice and has a substantial protective effect against WD-induced disruption of the endothelial barrier in mouse aortas and against the formation of atherosclerotic lesions in aortas of hypercholesterolemic LDLR^-/-^ mice. These data provide strong evidence that lipid-induced endothelial stiffening is a major contributor to vascular disease. Furthermore, mechanistically, we show that endothelial stiffening is differentially induced by specific LCFAs via GDI-1 inhibitory protein, demonstrating a novel mechanism of endothelial stiffening.

Early studies from our lab demonstrated that exposure to oxLDL results in significant stiffening of aortic ECs *in vitro*^36^, which we found to be a result not of an increase in endothelial cholesterol but of an incorporation of specific oxysterols^13,37^. In fact, cholesterol enrichment of aortic ECs via MβCD-cholesterol complex reversed oxLDL-induced stiffening, probably because of the exchange between cholesterol and oxysterols that induced the stiffening^13^. A receptor responsible for the uptake of oxLDL into HAECs and for the resulting stiffening was identified to be CD36, whereas another endothelial scavenger receptor LOX1^38^ had no role in oxLDL uptake^5,14^. We also found significant stiffening of aortic endothelium *in vivo*, in male mice fed WD for 4-6 weeks, which was abrogated by the global deletion of CD36 and was independent of CD36 in hematopoietic cells^5^. Our current study demonstrates that the stiffening effect critically depends on endothelial CD36.

Notably, endothelial stiffening is distinct from arterial stiffening, or stiffening of the vascular wall, a known predictor of cardiovascular disease. Arterial stiffening is a result primarily of the remodeling of the extracellular matrix^39^ and is typically observed after a prolonged WD^40,41^ or during ageing^42^. Earlier studies suggested that the pathological effect of arterial stiffening is at least in part due to its effect on the endothelium. Specifically, exposure of endothelial cells to a stiff substrate activates RhoA/ROCK cascade^1,3^, which multiple studies showed induces endothelial stiffening/contractility^8,14^ and barrier disruption^8,15^. Thus, it was proposed that stiffening of the vascular wall results in the stiffening of the endothelium via substrate-dependent activation of RhoA/ROCK, which in turn leads to the disruption of the endothelial barrier. Focusing on early stage of WD-induced vascular damage, our study shows, however, that in WD-fed mice endothelial stiffening and barrier disruption develop prior and independently of the arterial stiffening.

In terms of the functional significance of CD36-mediated endothelial stiffening, we demonstrate that endothelial CD36 plays a critical role in the endothelial barrier disruption and formation of atherosclerotic lesions. Previous studies found that deletion of CD36 may be both protective or disruptive for the maintenance of the barrier in a tissue-specific way: in the lung vasculature, deletion of CD36 was shown to be barrier protective in a mouse model of malaria infection^43^, whereas in small intestine and lymphatic endothelium, deletion of CD36 led to barrier disruption^27,44^. The mechanisms for these distinct effects are not clear. Our study in aortic endothelium shows that deletion of CD36 has no effect on the endothelial barrier under basal conditions and is protective in WD-fed animals, which corresponds to the changes in the endothelial elastic modulus. Given our findings that CD36-mediated uptake of oxLDL or saturated long chain fatty acids induce endothelial stiffening by increasing endothelial contractility, we suggest that this is a major mechanism by which CD36-mediated lipid uptake may induce barrier disruption. Taking these studies further to a hypercholesterolemic LDLR^-/-^ mouse model of atherosclerosis, we show that endothelial stiffening correlates with the severity of plaque formation when comparing males vs. females or as a result of downregulation of CD36. The role of CD36 in atherosclerosis was established previously: earlier studies showed that the global deletion of CD36 is atheroprotective^45^, which was attributed mainly to macrophages, and a recent study showing that EC-CD36^-/-^is also atheroprotective^22^, but the mechanism of the EC-CD36^-/-^ atheroprotection remained unclear. One possible explanation was that it is due to a global effect of decreased triglycerides in the serum but it is not clear whether a decrease in triglycerides without a significant decrease in LDL would be sufficient to have a significant atheroprotective effect. Our study suggests a specific mechanism by which CD36-mediated endothelial uptake of lipids may contribute to the initiation of plaque formation: altering endothelial biomechanical properties, which result in the disruption of the endothelial barrier, a hallmark of early stage inflammation.

Significant sex differences in the stiffening of aortic endothelium were observed in this study with males having more severe effects. In WD-fed WT mice, this difference correlated with lower levels of weight gain and smaller WD-induced increases in serum lipid levels. A difference in the lipid profiles and weight gain between males and females in response to WD indeed are well established, with most studies showing that females are relatively protected^18,21,22^, which can easily explain the sex difference in the endothelial stiffening. In terms of atherosclerosis development, previous studies were more controversial with some studies seeing more effects in males and some in females^46,47^. In our study, we consistently see more atherosclerotic plaques in male LDLR^-/-^ on WD than in female LDLR^-/-^, which corresponds to their respective differences in EC stiffening. Since abrogating endothelial stiffening in LDLR^-/-^ male mice has a strong protective effect against atherosclerosis, we propose that the sex differences in the severity of atherosclerosis in this model may also be a result of the differential stiffening effects.

This study also provides new insights into the mechanism of dyslipidemia-induced endothelial stiffening. Previously, we showed that endothelial stiffening is induced by oxLDL and its components^5,6,13,14,36,48^. Here, we establish that endothelial stiffening is also induced by LCFAs, which is significant because LCFAs are a major type of circulating lipids^49,50^. Furthermore, LCFAs have distinct effects on endothelial stiffening: saturated LCFA, palmitic and stearic acids, induce the stiffening effect, whereas non-saturated LCFA, oleic acid, does not. This corresponds to the known pro-inflammatory effects of palmitic acid and highlights the role of endothelial stiffening in lipid-induced inflammation. In terms of the signaling pathway responsible for the LCFA-induced endothelial stiffening, we show that similarly to oxLDL/oxysterols^14^, it is mediated by the ROCK activation. In contrast, activation of Src, a downstream target of CD36, does not contribute to the stiffening effect. A key question, however, is the mechanistic link between LCFA and RhoA activation. In this study, we address a hypothesis that LCFA may activate RhoA by interfering with its interaction with a known inhibitor of RhoA, Rho-GDI-1^35,51^. This hypothesis is based on the fact that RhoA interacts with GDI-1 via RhoA hydrophobic tail being inserted into a hydrophobic pocket of GDI-1^35^. Consistent with this idea, we have shown earlier that overexpression of Rho-GDI-1 but not endogenous inhibition of p115RhoGEF (via p115-RGS) inhibits oxLDL-induced RhoA activation^52^. Here, we show that expression of Rho-GDI-1 is essential for LCFA-induced endothelial stiffening, suggesting that uncoupling of RhoA from Rho-GDI-1 at its hydrophobic pocket underlies the mechanism of LCFA-induced RhoA activation and consequently endothelial stiffening. This constitutes a novel mechanism of lipid-mediated regulation of RhoA signaling and cellular stiffness.

Translationally, we propose that an early intervention to prevent WD-induced endothelial stiffening, barrier disruption and formation of atherosclerotic lesions may focus on inhibiting endothelial uptake of lipids rather than arterial stiffening. Our study suggests that endothelial CD36 can be a target to achieve this goal.

## NON-STANDARD ABBREVIATIONS AND SYNONYMS

AA: Ascending aorta or aortic arch
AFM: Atomic force microscopy
CD36: Cluster of differentiation 36 or fatty acid translocase
DA: Descending aorta
ECs: Endothelial cells
EBD: Evans blue dye
FAs: Fatty acids
HAECs: Human aortic endothelial cells
LCFA: Long-chain fatty acid
LFD: Low-fat diet
MβCD: Methyl-β cyclodextran
oxLDL: Oxidized low-density lipoprotein
PBS: Phosphate buffer solution
RT-qPCR: Real time-quantitative polymerase chain reaction
ROCK: Rho-associated protein kinase
Rho-GDI1: Rho guanosine dissociation inhibitor 1
SSO: Sulfo N-succinimidyl oleate
VE-Cadh: Vascular endothelial-cadherin
WD: Western diet
WT: Wild-type

## ACKNOWLEDGEMENTS

First we thank Prof Richard Minshall (UIC) for multiple insightful discussions and critical reading of the manuscript. We also thank Prof Kishore Wary (UIC) for the gift of Cdh5.CreER^T2^ mice. We also thank the UIC RRC Cardiovascular Core, specifically Director Dr Jiwang Chen, for his consultation and assistance with performing echocardiography. We also thank the UIC RRC Imaging Core, specifically Director Dr Peter Toth, for assistance with confocal microscopy imaging of mice aortas.

## HIGHLIGHTS

High fat diet-induced stiffening of the aortic endothelial layer associates with barrier disruption and is dependent on endothelial-specific expression of scavenger receptor CD36.

Prevention of aortic endothelial stiffening by downregulation of endothelial CD36 alleviates the formation of atherosclerotic lesions.

Endothelial stiffening is induced by saturated long-chain fatty acids through the activation of RhoA/ROCK signaling and is critically dependent on Rho-GDI1.

